# DUCK-Net: Automated deep learning segmentation of Ductular Reactions in murine liver injury captures multicellular niche dynamics from H&E morphology

**DOI:** 10.64898/2026.02.09.704844

**Authors:** Nathalie Feeley, Kai Williams, Daniel Field, Caitlin McCaffrey, Kyle Davies, Hugh Warden, Luke Boulter, Steve Thorn, Stephen J. Wigmore, Ewen M. Harrison, Stuart J. Forbes, Ian P. Tomlinson, Timothy J. Kendall, Rachel V. Guest

**Author notes:** **Address for correspondence:** Dr. Rachel V. Guest, Lab E3.07, Institute of Genetics and Cancer, University of Edinburgh, Crewe Road South, Edinburgh, United Kingdom EH4 2XU. **Author Contributions:** N.F.: Conceptualisation, methodology, software, validation, formal analysis, data curation, writing (original draft, review & editing), visualisation, project administration. K.W.: Conceptualisation, methodology, software, validation, formal analysis, data curation, writing (original draft, review & editing), visualisation. D.F.: Data curation. C.M.: Investigation, data curation. K.D.: Investigation. H.W.: Methodology, software. L.B.: Conceptualisation, resources. S.T.: Writing (review & editing). S.J.W.: Writing (review & editing). E.M.H.: Conceptualisation. S.J.F.: Conceptualisation, writing (review & editing). I.P.T: Conceptualisation, validation, supervision, funding acquisition. T.J.K.: Methodology, validation, writing (review & editing). R.V.G.: Conceptualisation, methodology, validation, formal analysis, investigation, resources, data curation, writing (original draft, review & editing), visualisation, supervision, project administration, funding acquisition. **Conflict of Interest Statement:** The authors have no conflicts of interest to declare. **Ethical Approval Statement:** All animal work included in this study was approved by the University of Edinburgh Animal Welfare and Ethics board (PPL Nos. PP3266462, PPD31D3D4) in accordance with the UK Animals (Scientific Procedures) Act 1986. See Materials & Methods for full details. **Patient Consent Statement:** Not applicable. **Permission to reproduce material from other sources:** Not applicable. **List of Abbreviations:** DR(s) Ductular Reactions(s); DUCK-Net Deep Understanding Convolutional Kernel Network; DDC 3,5-Diethoxycarbonyl-1,4-Dihydrocollidine; IHC Immunohistochemistry; *R*^2^ Coefficient of determination; H&E Haematoxylin and eosin; PSC Primary Sclerosing Cholangitis; PBC Primary Biliary Cirrhosis; BEC(s) Biliary Epithelial Cells; HPC(s) Hepatic Progenitor Cells; ECM Extracellular matrix; MASLD Metabolic Dysfunction-Associated Liver Disease; CNN Convolutional Neural Network; PBS Phosphate-buffered saline; KRT19 Cytokeratin 19; SOX9 SRY-Box Transcription Factor 9; α-SMA Alpha-smooth muscle actin; DAB 3,3-diaminobenzidine; PSR Picrosirius red; RGB Red-Green-Blue; HED Haematoxylin-eosin-DAB; CIRSTA Cholestatic Injury and Repair Spatio-Temporal Atlas; LPLC Liver Progenitor Like Cell(s); Chol Cholangiocyte; ROC Receiver Operating Characteristic; AUC Area Under the Curve; CI Confidence Interval; RMSE Root mean square error; DTW Dynamic Time Warping; WSI Whole slide image; SD Standard deviation; SE Standard error; MAP Mean pixel accuracy; DNB DNA nanoball.

## Abstract

Ductular Reactions (DRs) are dynamic and complex multicellular responses that occur as a result of various hepatic injuries. Precise identification and quantification of the extent of DRs is a cornerstone of pre-clinical modelling of liver disease, with links to inflammation, fibrosis, regeneration, and disease severity. Here, we apply a deep learning model, Deep Understanding Convolutional Kernel or ‘DUCK-Net’, to the automated detection and segmentation of DRs in whole-slide histopathological images of murine models of liver damage. Following annotation of a training dataset by a specialist liver histopathologist, we demonstrate accelerated performance and accurate detection, achieving a mean Dice coefficient (model-expert segmentation overlap) of 85.4% and a specificity of 98%, indicating minimal false positives. Evaluation of model validity and utility was achieved with a histological time course of cholestatic injury and recovery using 3,5-Diethoxycarbonyl-1,4-Dihydrocollidine diet (DDC) in mice. When assessed against a multiple linear regression model incorporating core epithelial and stromal components of the DR as quantified using IHC, DUCK-Net predicted the spatiotemporal response to injury and repair/resolution with a coefficient of determination (*R*^2^) of 0.88. Moreover, DUCK-Net kinetics strongly correlated with published spatial transcriptomic (Stereo-seq) analysis of the DDC model, demonstrating that H&E-based segmentation captures molecular DR dynamics comparable to, or exceeding that of individual IHC markers without the need for immunostaining. DUCK-Net provides a novel and accessible platform for rapid, accurate histological quantification of liver injury reflective of the matrix-rich, multicellular regenerative niche observed in DRs.

## Introduction

Cholestatic liver diseases such as primary sclerosing cholangitis (PSC), primary biliary cirrhosis (PBC), and other cholangiopathies are associated with the expansion of periportal ductular cells, known histologically as the Ductular Reaction (DR) ^1,2^. In addition to the proliferation and/or hyperplasia of biliary epithelial cells (BECs) and Hepatic Progenitor Cells (HPCs), DRs are characterised by fibrogenesis and infiltration of stromal and inflammatory cells including macrophages into the periportal regions ^3^. DR expansion has been shown to be accompanied by angiogenesis ^4^ and, in certain contexts, transdifferentiation of hepatocytes ^5,6^. Deposition of extracellular matrix (ECM), in particular laminin, within the DR maintains a local ‘niche’ to determine cell fate of HPCs and modify the cellular response according to injury aetiology ^7^. Further to the well-established association with biliary injury, DRs can also be observed in the context of severe hepatocyte damage including alcoholic hepatitis and Metabolic Dysfunction-Associated Liver Disease (MASLD) and the extent of DR can act as a histological correlate of disease severity and patient mortality ^3,8,9^.

Dietary 3,5-Diethoxycarbonyl-1,4-dihydro-collidine (DDC) in mice recapitulates the effects of human cholestatic disease by inducing periportal injury through induction of interlobular biliary obstruction with protoporphyrin plugs. Toxic accumulation of bile acids activates BECs and initiates periportal inflammation and fibrosis as well as secondary hepatocyte damage as evidenced by raised serum transaminases and parenchymal necrosis ^10,11^. BECs up-regulate genes that trigger recruitment and differentiation of macrophages, further stimulating BEC proliferation, as well as inducing HPC differentiation into BECs and dampening the proliferation of periportal hepatocytes ^12^. The consistent induction of periportal DRs by DDC within weeks of feeding and resultant recovery following dietary withdrawal has made it a frequently used model of liver injury and regeneration and its spatiotemporal dynamics have been widely studied ^12^.

The quantification of DRs within histopathological slides of murine liver tissue is a complex and time-consuming process subject to inter-observer variability. A manual approach to annotation requires histopathology experience, creates a bottleneck in research workflow, and introduces inconsistencies in murine studies. Recent advances in deep learning models have facilitated the automation of image analysis in medical research. Computer vision (the interpretation of visual information by machines) and in particular image segmentation have showcased the ability to identify complex structures in medical images. Models such as U-Net, Mask R-CNN, and DeepLab have been developed specifically for the detection of nuclei, cells, and complex tissue boundaries or other multi-scale features within histological slides ^13–15^. DUCK-Net (Deep Understanding Convolutional Kernel), a specialised neural network architecture has been developed for use in the segmentation of human colonic polyps, enabling the automated detection and location of polyp borders. This convolutional neural network (CNN) combines the encoder-decoder structure of U-Net, an existing deep learning architecture, and a custom-designed convolutional block with residual down sampling. DUCK-Net uses six variations of convolutional blocks in parallel, allowing the network to adaptively select the most effective features during training. Low-level detail is preserved by the addition of a secondary U-Net downscaling layer that bypasses image processing ^16^. This improves on previous models through the accurate prediction of border location, with the ability to utilise initial information at each resolution level in the encoder segment, allowing the model to learn effectively from small datasets.

This multi-resolution approach is ideally suited for histological images where cellular structures vary in size and appearance, allowing complex segmentation of heterogenous multicellular regions such as those seen in liver injury, and detection of subtle differences between cell populations. The use of data augmentation allows DUCK-Net to handle variability in tissue staining and preparation, common challenges in histological analysis of slides of liver injury. In this study, we have implemented DUCK-Net for the production of annotations of DRs present in murine livers treated with DDC diet and evaluate its predictive accuracy to the ground truth annotation. To provide orthogonal validation of H&E segmentation using DUCK-Net to accurately capture the complex multi-component DR, we present correlation analyses with core classical IHC markers as well as molecular data using a previously published, publicly available spatial transcriptomics dataset (Stereo-seq) from an independent DDC study ^12^.

## Materials and Methods

### Animal experiments

C57BL/6 mice were bred in-house at the University of Edinburgh, UK and housed in pathogen-free conditions. All experimental protocols were approved by the University of Edinburgh Animal Welfare and Ethics Board in accordance with UK Home Office regulations. At 6 to 8 weeks old, female mice were fed 0.1% 3,5-Diethoxycarbonyl-1,4-dihydro-collidine for up to 21 days and placed back onto standard chow for the recovery model. Four mice were culled at 6 timepoints. Mice fed normal chow were used as a baseline control group. Sample size was calculated *a priori* based on published Krt19+ area quantification data from Fickert et al. in the DDC model^17^. Assuming group means of 5 (control), 8 (peak injury), and 6 (recovery) arbitrary units with a pooled standard deviation of 1, the effect size (Cohen’s f) was 1.25. Using one-way ANOVA with α = 0.05 and 80% power, n = 4 mice per group were deemed to be required, providing 91% power to detect significant differences across the injury-recovery time course.

### Immunohistochemistry and slide analysis

Mice were culled via a Schedule 1 procedure and livers perfused with PBS via the Inferior Vena Cava. Livers were fixed overnight in 10% buffered formalin before paraffin embedding, cutting 4μm sections and staining with haematoxylin and eosin. Sections were deparaffinised in xylene and rehydrated through serial alcohols. The following antibodies were used: cytokeratin 19 (TROMA-III) (AB 21335570 Developmental Studies Hybridoma Bank), SRY-Box Transcription Factor 9 (SOX9) (Sigma HPA001758), and α-smooth muscle actin (α-SMA) (Sigma A2547). Heat-induced epitope retrieval was performed through immersion in Tris-EDTA buffer (10mM Tris, 1mM EDTA, pH9.0) for 10 minutes before staining. Immunoreactivity was visualised using 3,3-diaminobenzidine (DAB) substrate solution with hydrogen peroxide in phosphate-buffered saline (PBS) according to the manufacturer’s instructions (Dako). To stain pan-collagen fibres, sections were incubated with picrosirius red solution (0.1% Sirius Red in saturated picric acid) for 60 minutes at room temperature followed by rinsing with 0.5% acetic acid solution. Sections were digitised using a Hamamatsu NanoZoomer S360MD whole slide imaging scanner and manual annotations made using QuPath open-source software (version 0.3.2)^18^.

### Data preparation and annotation

Training and validation data were derived from H&E-stained liver sections from mice following 21 days of DDC diet. The primary training dataset consisted of a whole slide image (118,784 x 107,520 pixels) selected for abundant DR, with a focal region of blur excluded to prevent ambiguous training signals. To capture and account for heterogeneity in DR and staining characteristics, regions from 50 further slides were selected to represent diverse staining intensities and tissue morphologies. All annotations were performed by a clinical pathologist, who annotated 1,596 polygon regions across training and validation images. Annotation followed a standardised protocol, with biliary populations identified based on morphological characteristics ^19^. For model testing, 10 randomly sampled regions (21,216 × 20,840 pixels) were selected from unique 21-day DDC diet sections not included in training or validation, representing high DR prevalence.

### Model Architecture, Validation and Testing

Full details on model architecture and training are provided in the Supplementary materials. Code for the DUCK-Net model and analysis have been made publicly available at: https://github.com/WhatsMyPurpose/ducknet-dr-segmentation. Model validation was performed at the end of each training epoch using full-image inference on the validation set. Final model performance was evaluated on the held-out test set of 10 annotated samples, which were not used during training or hyperparameter selection. Final statistics for annotated pixels and tissue coverage in the training, validation and test datasets are reported in Supplementary Table S1.

### Datasets

Raw spatial transcriptomic data (Stereo-seq and scRNA-seq) from Wu et al.^12^ were downloaded from the CNGB Nucleotide Sequence Archive (accession number CNP0003447; https://db.cngb.org/search/?q=CNP0003447). Processed H5AD files containing spatial gene expression profiles were accessed through the Cholestatic Injury and Repair Spatio-Temporal Atlas (CIRSTA) database (https://db.cngb.org/stomics/cirsta/). Liver Progenitor Like Cells (LPLC) and Cholangiocyte (Chol) domain scores representing the spatial transcriptomic signatures of DRs were extracted for each timepoint (days 0, 8, 17 during DDC injury; days 19, 24, 38 during recovery) using the Python *anndata* library (v0.9.2, Python) accessed via the *R reticulate* package (v1.28, R).

### Statistical Analysis

Assessment of the performance of the DUCK-Net model against ground truth clinical annotations was evaluated via multiple metrics including Dice coefficient, volumetric overlap, precision, recall (sensitivity), specificity, accuracy, and area under the receiver operator curve (ROC). To determine whether DUCK-Net segmentation could be explained by individual IHC markers over the DDC time course, a multiple linear regression model was selected using DUCK-Net coverage as the dependent variable and individual IHC markers of DR components as the independent variables (KRT19, SOX9, α-SMA, picrosirius red). Model fit was assessed using R^2^ and F statistics. Receiver Operating Characteristics (ROC) analysis was performed to evaluate the discriminatory ability of DUCK-Net and conventional IHC markers for classifying injury states in the DDC model (Supplementary Methods).

To enable comparison between published spatial transcriptomic data and our DUCK-Net DDC time course, (days 0,7,14,21 injury; days 28, 35 recovery), both the Wu et al. dataset and our DDC time course were interpolated to common timepoints (0, 7, 14, 21, 28, 35 days) using linear interpolation (approx() function in R, rule=2). Interpolation was validated by comparison with cubic spline and LOESS methods. To account for different experimental protocols (21-day vs. 17-day DDC treatment), time courses were normalised to % of protocol duration to demonstrate universal kinetics independent of absolute study duration. Linear regression was performed separately for injury and recovery phases to compare kinetic slopes. Pearson correlation coefficients were calculated between LPLC domain scores and each marker (DUCK-Net, KRT19, SOX9, α-SMA, PSR) using interpolated time courses. Statistical significance was assessed using cor.test() function with two-tailed tests.

## Results

To evaluate the ability of DUCK-Net to quantify DRs from H&E morphology, we first assessed model performance against expert pathologist annotations on an independent test set of 10 diverse mouse liver sections selected to represent varying DR severities and tissue morphologies following 21 days DDC dietary injury. Visual comparison demonstrated apparent close spatial concordance between DUCK-Net predictions and pathologist-annotated DR regions on both the initial training slide (Figure 1A-B) and the test set (Figure 1C), including accurate delineation around regions of small and large ducts, anastomosing ductular networks, and interface regions between portal tracts and parenchyma (Figure 1C). Quantitative evaluation across the test set demonstrated robust performance (Supplementary Table S2), achieving a Dice coefficient of 0.85 ± 0.05 (Jaccard index 0.74), with consistency across diverse tissue appearances and staining variations (Figure 1D). Training curves demonstrated effective learning of the model, with validation loss decreasing rapidly initially before plateauing, indicating successful model convergence without overfitting over 88 epochs of training (Figure 1E). The model achieved robust performance metrics across the test set (Figure 1F; individual sample data in Supplementary Table S2): precision 0.84 ± 0.07, recall 0.88 ± 0.08, mean pixel accuracy (MAP) 0.96 ± 0.01, and notably specificity was high (0.98 ± 0.01), indicating excellent discrimination against false positive detection.

**Figure 1.**
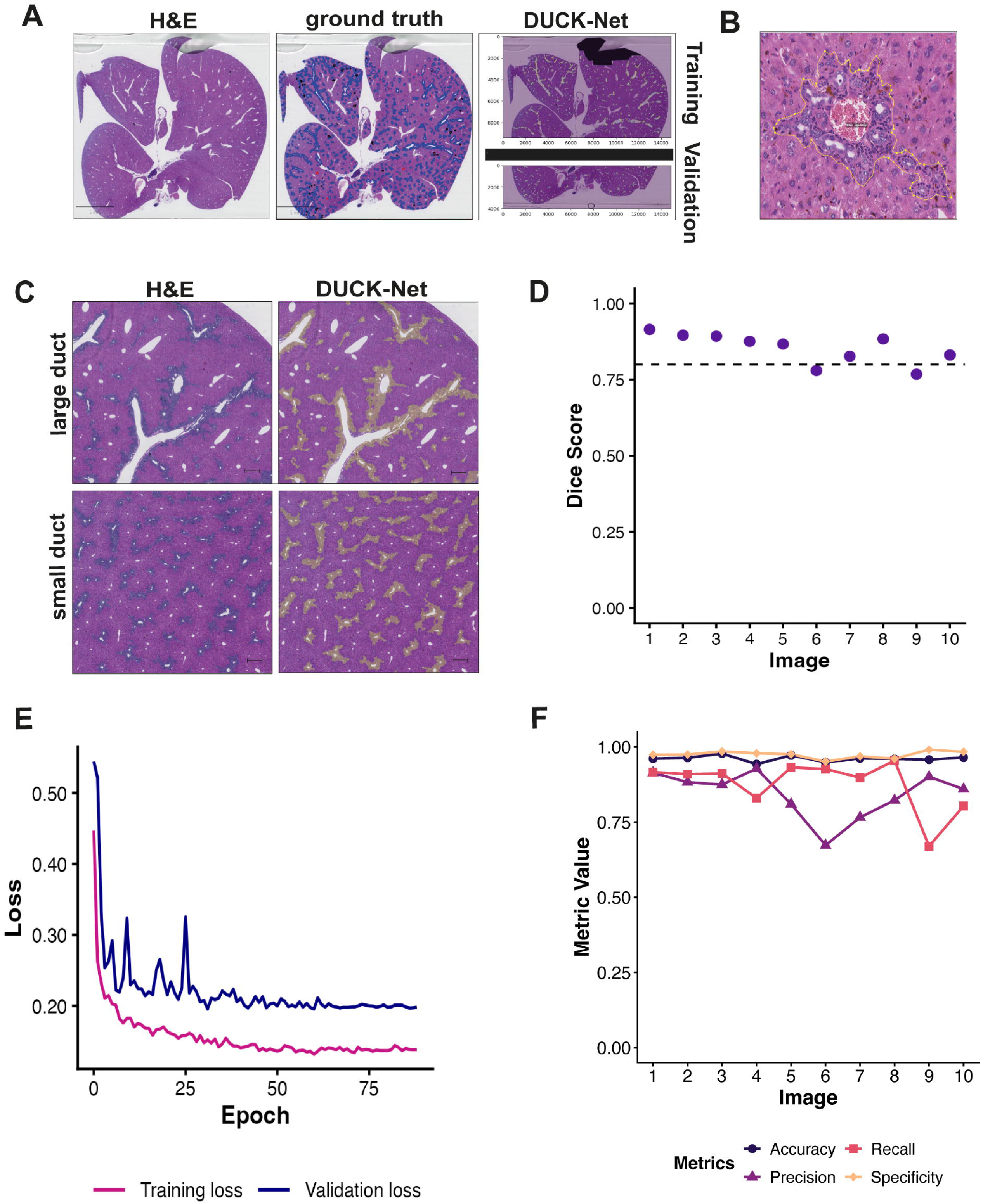
DUCK-Net accurately quantifies ductular reaction from H&E morphology in mouse liver tissue following DDC dietary injury. (A) Representative mouse whole liver section (H&E) following 21 days DDC treatment with expert pathologist annotations (ground truth) (blue overlay) and DUCK-net segmentation (yellow overlay) in training and validation regions of initial training slide. Artefact masking and eosin channel enhancement within training dataset. Scale bars 5mm. (B) High magnification view of DR region as annotated (yellow) by clinical liver histopathologist, capturing epithelial ductular structures and surrounding reactive stroma. Scale bar: 100μm. (C) Representative H&E and DUCK-Net predictions from test set showing segmentation across diverse morphologies including large ducts (top) and small ducts (bottom) and interface regions. Scale bars: 200μm. (D) Dice coefficients for 10 test samples (mean 0.85 ± 0.05) demonstrating consistent performance across varying DR severities. Dashed line indicates mean value. (E) Training and validation loss curves across 88 epochs. Smooth convergence indicates effective learning without overfitting. **(F)** DUCK-Net performance across 10 test samples including precision, recall (sensitivity), specificity and accuracy.

To assess whether DUCK-Net segmentation was able to faithfully map the appearance, progression and resolution of DRs, DUCK-Net coverage was quantified on H&E-stained sections from a DDC-induced model of liver injury and recovery (0.1% DDC diet, n=4 per time point) (Figure 2A). A panel of core established immunohistochemistry markers were selected to characterise the spatiotemporal dynamics of DRs: KRT19 and SOX9 for ductular epithelium and α-SMA and picrosirius red for the associated stromal response (Figure 2B). DUCK-Net coverage increased progressively during DDC injury, peaking at day 21 before declining during the recovery phase (Figure 2C). This temporal pattern closely mirrored the dynamics of all 4 IHC markers. Notably, DUCK-Net detected the onset of ductular reaction at day 7 and captured the expected lag in resolution during recovery, consistent with the known biology of DR regression.

**Figure 2.**
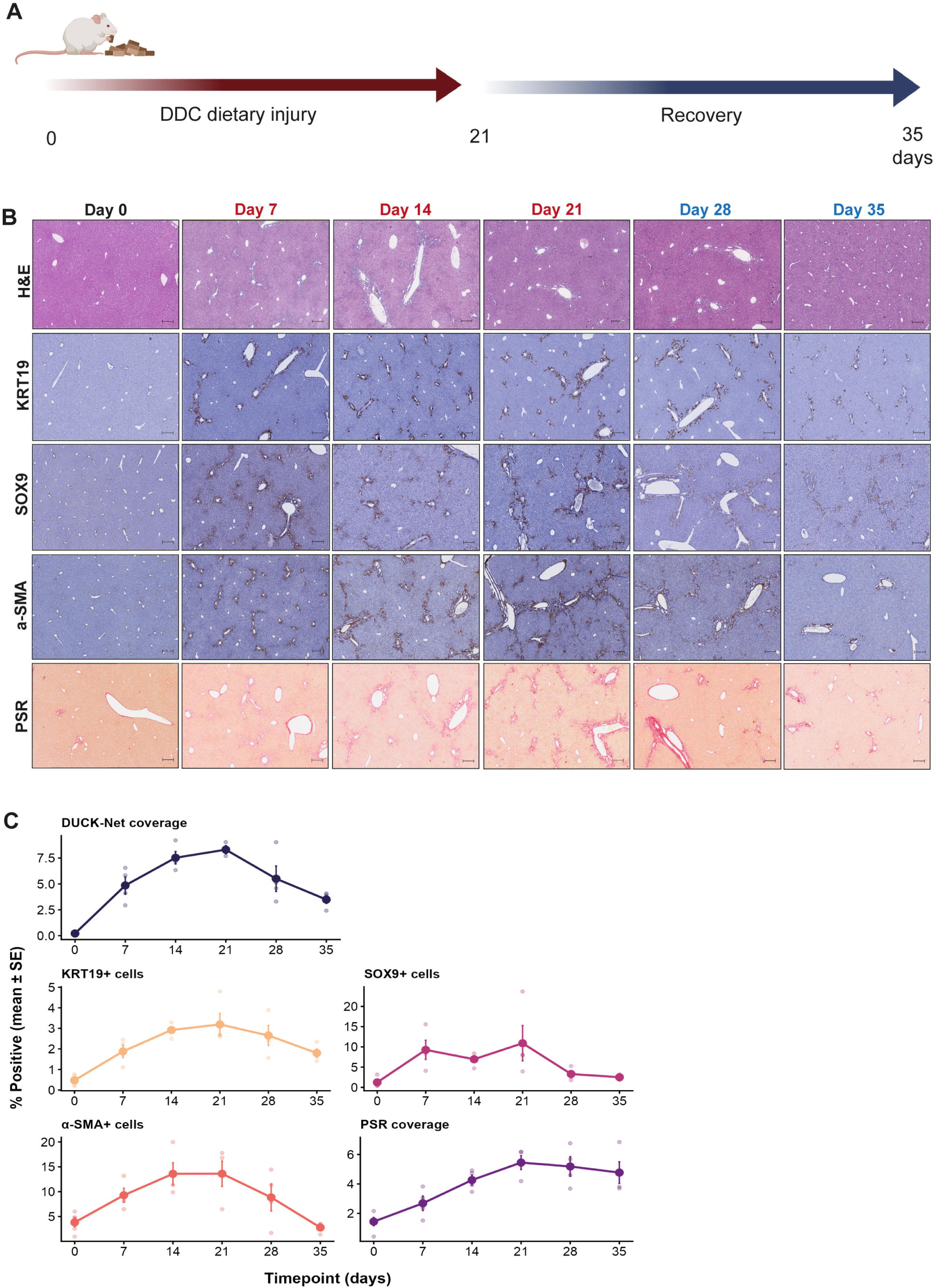
DUCK-Net and IHC marker quantification across DDC-induced liver injury and recovery. (A) Schematic of experimental timeline. Mice were fed 0.1% 3,5-Diethoxycarbonyl-1,4-dihydro-collidin (DDC) diet for up to 21 days followed by 14 days of recovery on normal chow (n=4 per timepoint). (B) Representative images of liver sections stained with H&E or immunohistochemical markers of ductular reaction across the DDC time course: cytokeratin 19 (KRT19), SRY-Box Transcription Factor 9 (SOX9), alpha smooth muscle actin (α-SMA), picrosirius red (PSR). Scale bars represent 200 μm. (C) Quantification of DUCK-Net coverage and IHC markers across the time course. DUCK-Net coverage was expressed as percentage coverage on H&E sections. IHC markers were quantified as percentage positive cells (KRT19, SOX9, α-SMA) or percentage area (PSR). Markers show a progressive increase during injury (days 0-21) and decline during recovery (days 28-35). Data are shown as mean ± SE (n = 4 per time point).

In order to investigate whether DUCK-Net coverage is able to fully capture the DR niche, and representation of both epithelia and stroma, a multiple linear regression model was constructed to model DUCK-Net variance as a function of core DR components (KRT19, SOX9, α-SMA, PSR) across the DDC time course (Table 1). Strong correlations were observed between DUCK-Net coverage and all IHC markers individually (KRT19 *r* = 0.90, *p* < 0.0001; α-SMA *r* = 0.69, *p* = 0.0005; PSR *r* = 0.52, *p* = 0.015; and SOX9 *r* = 0.51, *p* = 0.019) (Figure 3A). Model assumptions were validated by examining pairwise relationships between all variables using scatterplot matrices and residual diagnostics (Supplementary Figure 1A-B). Strong agreement was observed between IHC-predicted and actual DUCK-Net values across individual samples from all timepoints (*R*^2^ = 0.88, RMSE = 1.0) (Figure 3B) and DUCK-Net closely tracked predicted values throughout injury and recovery, with the model explaining a high proportion of the variance in DUCK-Net coverage across the time course (*R*^2^ = 0.88, *F*_4,16_ = 30.46, *p* = 2.66×10^-7^) (Figure 3C). Analysis of individual marker contributions revealed that KRT19 was the dominant predictor (β = 2.14, *p* = 0.00016), with SOX9 also contributing significantly (β = 0.20, *p* = 0.028). The stromal markers α-SMA and PSR did not reach significance as independent predictors in the multivariate model (Figure 3D). Despite the observed moderate correlation between KRT19 and SOX9 (*r* = 0.33, *p* = 0.15), both epithelial markers remained independent predictors, suggesting they capture at least partially distinct populations within the ductular epithelium (Figure 3E). Twelve percent of DUCK-Net variance was not explained by the model and is likely to reflect additional morphological features not captured by this core IHC panel such as inflammatory infiltrate, tissue architecture, or cellular morphology. This was maximal at the point of peak injury (days 14–21), when DUCK-Net tended to exceed predicted values (Figure 3F).

**Figure 3.**
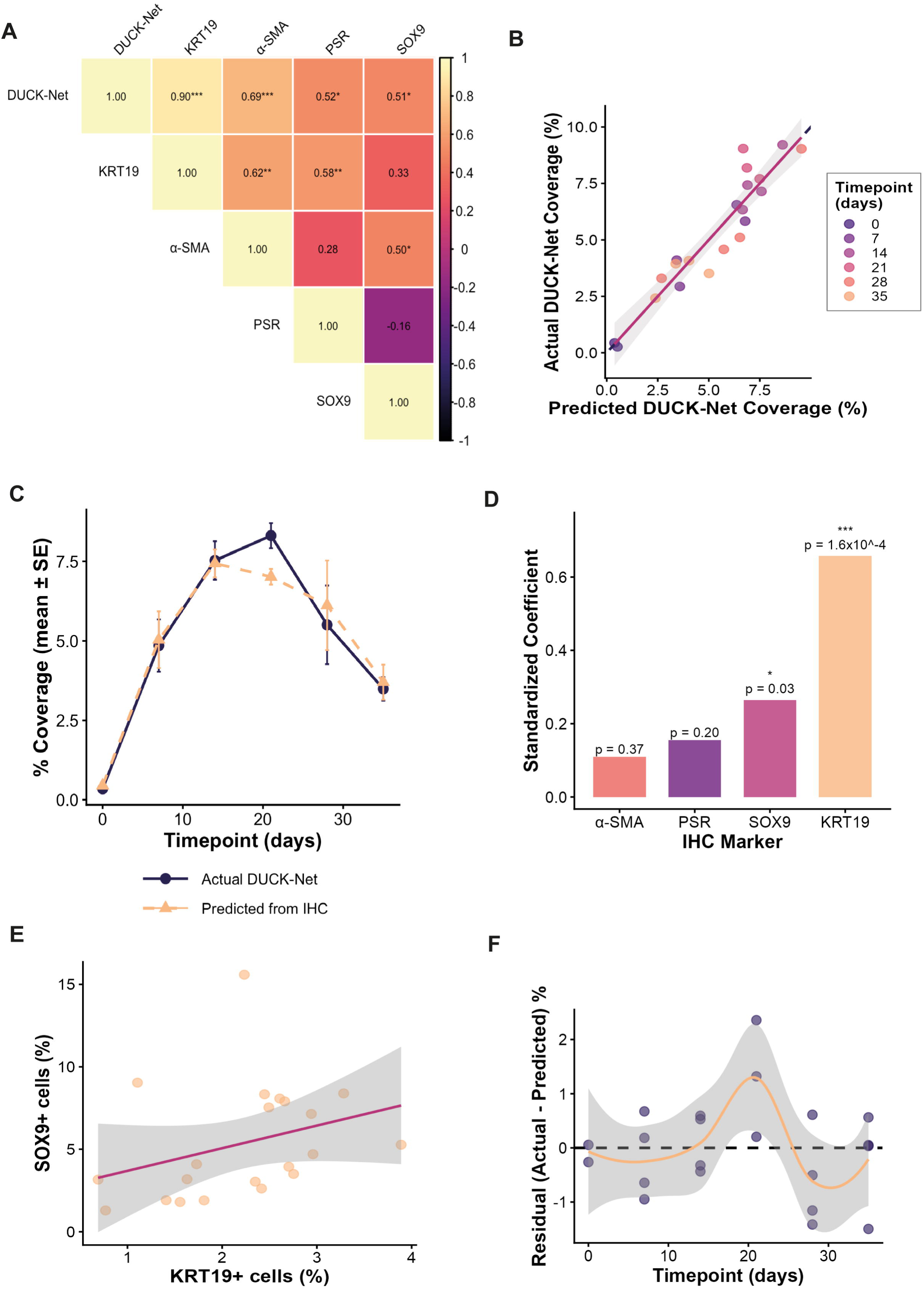
DUCK-Net captures the DR niche as characterised by IHC markers across a time course of DDC injury and recovery. (A) Correlation matrix showing Pearson correlation co-efficients between DUCK-Net coverage and individual IHC markers. Colour intensity reflects correlation strength (scale bar, right). Significance levels \**p* < 0.05, \*\**p* < 0.01, \*\*\**p* < 0.001. (B) Scatter plot of predicted versus actual DUCK-Net coverage for individual samples. Each point represents one sample, coloured by timepoint (days). Dashed line indicates perfect agreement (y=x); orange line shows linear regression fit with 95% confidence interval. *R*^2^ = 0.88, RMSE = 1.0. (C) DUCK-Net coverage (actual, blue) versus IHC predicted values (orange, dashed) across the DDC injury and recovery time course. Data shown as mean ±SE (n=4 per timepoint). Model: *R*^2^=0.88, *F*_4,16_=30.46, *p*=2.66×10^-7^. (D) Standardised regression coefficients showing relative contribution of each IHC marker to DUCK-Net prediction. Significance levels: ns not significant, \**p* < 0.05, \*\*\**p* < 0.001. (E) Scatter plot of SOX9 versus KRT19 expression, demonstrating moderate correlation between epithelial markers (*r* = 0.33, *p*=0.15). Line shows linear regression fit with 95% confidence interval. (F) Model residuals (actual – predicted DUCK-Net coverage) across the time course. Dashed line indicates zero residual; orange line shows LOESS smoothed trend with 95% confidence interval. Positive residuals at days 14-21 indicate DUCK-Net exceeds predicted values at peak injury.

**Table 1.** Multiple Linear Regression Coefficients (DUCK-Net versus IHC markers). Model fit statistics from multiple linear regression analysis modelling DUCK-Net coverage as a function of core DR components across DDC injury and recovery time course. Predictor variables comprised epithelial markers (KRT19, SOX9) and stromal markers (α-SMA, PSR). Beta, unstandardised regression coefficient; SE, standard error; Standardised Beta, regression coefficient normalised to allow comparison of relative predictor contributions; CI 95% confidence interval for unstandardised Beta. KRT19 and SOX9 were significant independent predictors of DUCK-Net coverage.

| Predictor | Beta | SE | Standardised<br>Beta | CI 95<br>Lower | CI 95<br>Upper | T value | P value |
| --- | --- | --- | --- | --- | --- | --- | --- |
| (Intercept) | -2.08 | 0.80 | - | -3.78 | -0.37 | -2.58 | 0.0201 |
| KRT19 | 2.14 | 0.44 | 0.66 | 1.22 | 3.06 | 4.91 | <0.001 |
| α-SMA | 0.05 | 0.06 | 0.11 | -0.07 | 0.17 | 0.93 | 0.3673 |
| PSR | 0.27 | 0.20 | 0.16 | -0.16 | 0.69 | 1.33 | 0.2027 |
| SOX9 | 0.20 | 0.08 | 0.26 | 0.02 | 0.38 | 2.41 | 0.0283 |

To evaluate the discriminatory performance of DUCK-Net relative to conventional IHC markers, ROC analysis was performed, assigning clinically relevant categories (uninjured liver day 0; any injury day 7–35; peak injury day 14–21; early injury day 0–7 and recovery day 21–35) (Figure 4A, Supplementary Table S3). For distinguishing control from injured liver, DUCK-Net achieved perfect discrimination (AUC = 1.00), equivalent to KRT19 (AUC = 1.00) and outperforming α-SMA (AUC = 0.87). Similarly, for identifying peak versus early injury, DUCK-Net demonstrated excellent performance (AUC = 0.98, comparable to KRT19 (AUC = 1.00) and PSR (AUC = 0.98), whilst SOX9 showed no discriminatory ability (AUC = 0.50). Interestingly, in distinguishing recovery from any injury, SOX9 demonstrated superior discrimination (AUC = 0.97) followed by α-SMA (AUC = 0.82) and DUCK-Net (AUC = 0.81), whilst KRT19 and PSR performed poorly (AUC = 0.66 for both), reflecting the distinct biological processes captured by each marker.

**Figure 4.**
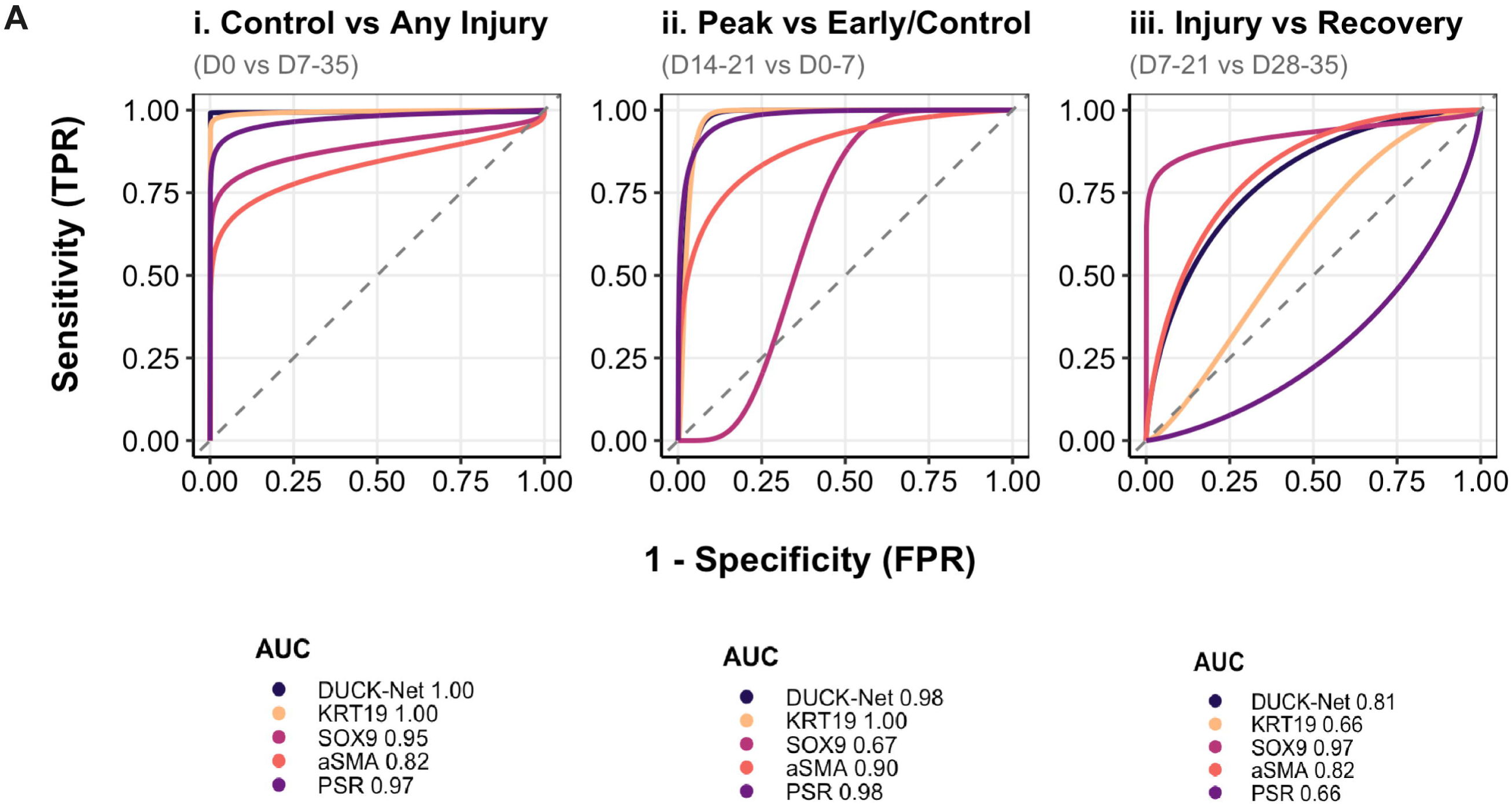
Discriminatory performance of DUCK-Net in identification of liver injury and recovery. (A) Receiver Operating Curves (ROC) comparing the discriminatory performance of DUCK-Net (blue) against conventional IHC markers (KRT19, SOX9, α-SMA and PSR). i. control versus any injury (day 0 vs day 7 – 35). ii. Peak injury versus early/control timepoints (day 14 – 21 vs day 0 – 7) and iii. active injury versus recovery phase (day 7 – 21 vs day 28 – 35). AUC values are shown below individual plots. Dashed diagonal line indicates performance of a random classifier (AUC = 0.50). Curves were smoothed with binormal fitting for visualisation only; all AUC values were calculated from empirical data. n = 4 at each time point.

Several comprehensive molecular characterisations of the DR niche using single cell resolution techniques have been described, including DDC-induced cholestatic liver injury^23,24^. These have included studies using spatial enhanced resolution omics sequencing (Stereo-seq), a DNA nanoball (DNB) method that applies high resolution gene expression data to a wide spatial area, ensuring signals from rare cell populations are not diluted by surrounding common cell types and allowing characterisation of regional variations across tissues^12^. To investigate whether DUCK-Net has potential to capture the molecular dynamics of the DDC damage response, publicly available raw spatial transcriptomic (Stereo-seq) and single-cell RNA-seq data as published by Wu et al. in *Nature Genetics* ^12^ were accessed from the CNGB Nucleotide Sequence Archive through the CIRSTA interactive database. Time course alignment was conducted using linear interpolation and normalised to protocol duration. The proportions of Cholangiocyte (Chol) *(Krt19^+^, Krt7^+^)* and Liver Progenitor Like Cell (LPLC) (*Spp1^+^*, *Sox9^+^*, *Krt19^−^*, *Krt7^−^*) spatial domains were extracted at each timepoint and compared with DUCK-Net coverage (Supplementary Table S4). DUCK-Net and LPLC domains shared similar kinetic trajectories during injury and recovery phases, both peaking at the point of DDC withdrawal and exhibiting progressive recovery but incomplete resolution at the end of the model (Figure 5A). In contrast, the Chol domain showed little dynamic change throughout the time course, with approximately 9-fold smaller magnitude of response compared to LPLC.

**Figure 5.**
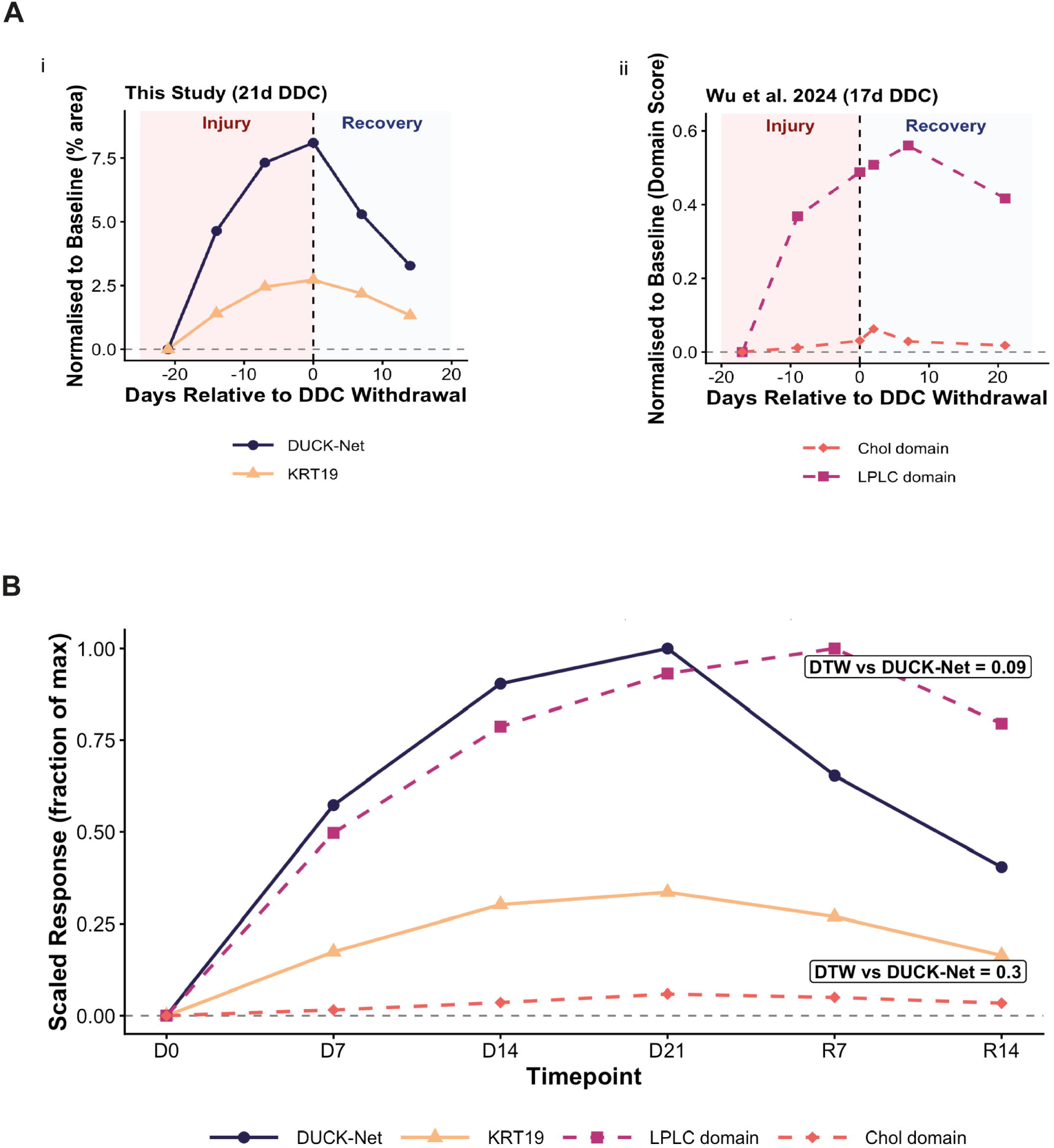
DUCK-Net correlates with spatial transcriptomics-defined liver progenitor cell domains from an independent dataset. (A) Universal ductular reaction kinetics across the 2 studies. DUCK-Net coverage and KRT19 quantification were normalised to protocol duration and compared with LPLC (Liver Progenitor Like Cell) and Cholangiocyte (Chol) domain coverage from spatial transcriptomic (Stereo-seq) data as published by Wu et al. (Nature Genetics 2024) accessed from the CNGB Nucleotide Sequence Archive. Timepoints were expressed as days relative to withdrawal of DDC diet. Values were normalised to baseline expression within each dataset. Linear interpolation was applied to estimate values at equivalent timepoints between studies. Datasets show concordant kinetics between DUCK-Net and LPLC domains, with progressive increase during injury and decline during recovery, demonstrating DR dynamics are conserved across experimental models and quantification methods. Shaded regions indicate injury phase (pale red) and recovery phase (pale blue). (B) Scaled trajectory comparison of DUCK-Net and KRT19 IHC quantification with LPLC and Chol spatial domains from CIRSTA (Wu et al., 2024). Data were normalised as change from D0 baseline and scaled within unit types. DUCK-Net (blue) and LPLC (pink) domain trajectories show closely matched kinetics, both peaking at the end of injury and declining during recovery. The Chol domain (dark orange) shows minimal dynamic change (approx. 9-fold smaller response than LPLC (light orange)). Dynamic Time Warping (DTW) distance from DUCK-Net is shown for each CIRSTA domain; lower values indicate greater trajectory similarity. Solid lines = this study; dashed lines = CIRSTA data.

To quantify trajectory similarity while accounting for differences in response magnitude, Dynamic Time Warping (DTW) was selected to enable quantitative comparison of trajectories in the two models using data normalised to baseline. DTW measures the similarity between two temporal sequences by computing the minimum cumulative distance required to establish the optimal alignment between them, and yields an output distance metric where lower values indicate more similar trajectories ^25–27^. DTW analysis revealed that DUCK-Net trajectories were 3.2-fold more similar to the LPLC domain (DTW distance = 0.091) than to the Chol domain (DTW distance = 0.295) (Figure 5B), (Table 2).

**Table 2.**
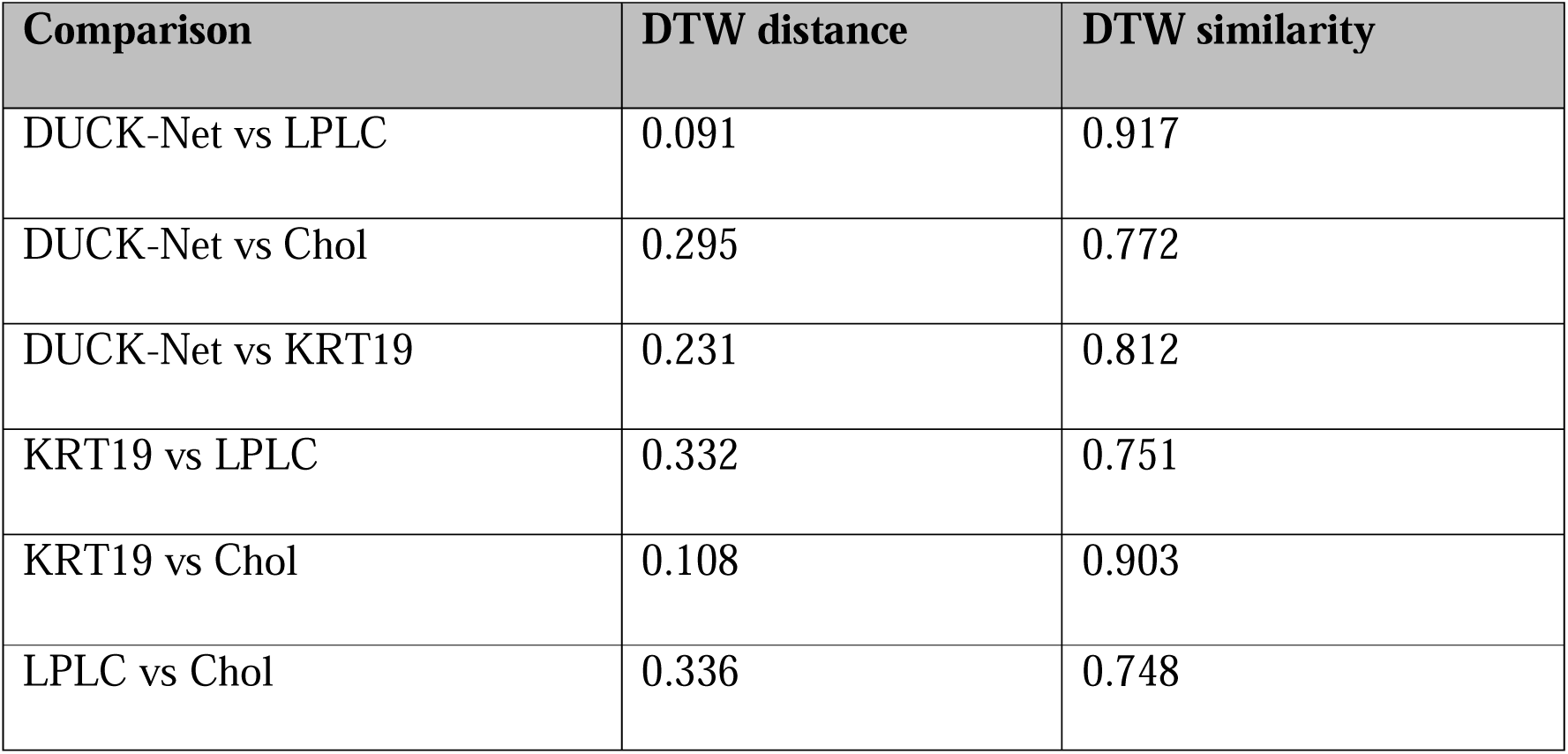
Dynamic Time Warping Analysis of DUCK-Net and CIRSTA Spatial Domains. Pairwise Dynamic Time Warping (DTW) analysis comparing temporal trajectories of DUCK-Net coverage and IHC marker (KRT19) quantification over the DDC injury/recovery time course with Liver Progenitor Like Cell (LPLC) and Cholangiocyte (Chol) spatial domain proportions from published Stereo-seq data (Wu et al., *Nature Genetics* 2024). DTW distance represents minimum cumulative distance required for optimal alignment between two temporal sequences; lower values indicate greater trajectory similarity. DTW similarity was calculated as 1/(1+DTW distance). All time courses were normalised to baseline (day 0) values. DUCK-Net trajectories showed strongest alignment with LPLC domains (DTW distance 0.091)

## Discussion

The development of deep learning algorithms to accurately and rapidly analyse medical images has generated considerable interest in the diagnosis, management and understanding of an array of disease states ^28^. Application of these tools to H&E-stained tissues has opened up new possibilities for prediction of molecular properties from morphological features, such as microsatellite instability and even patterns beyond human visual perception ^29^. Computer vision, specifically convolutional neural network (CNN) architectures, have been widely adopted as the models of choice for the segmentation of digital images, especially analysis of digital histopathology and H&E-stained whole slide images (WSI) ^30^. These have included architectures capable of creating highly detailed segmentation maps using limited training samples with extremely fast learning capability ^31^. DUCK-Net builds on such models, using a convolutional block to improve accuracy of prediction of complex or irregular border locations and residual down-sampling to utilise image information at each resolution level, keeping the original field of view alongside the processed input image. This prevents loss of multi-scale information to improve small target detection ^16^.

We hypothesised that the architectural features of DUCK-Net make it particularly well-suited to the detection of DRs in mouse liver injury models. DRs frequently do not exhibit sharp, well-defined boundaries and the interface regions between portal tracts and the parenchyma can be complex and irregular. Furthermore, DRs contain heterogenous histological structures which simultaneously span multiple scales: small individual cells including HPCs and ductules, large anastomosing ductular networks as well as the surrounding stroma and ECM, all exhibit different H&E staining characteristics. The parallel convolutional blocks used by DUCK-Net allow the network to adaptively select the most appropriate feature strategy for different tissue components. Given the well-documented spectrum of histopathological phenotypes across disease pathologies, various published classification systems, and challenges in annotation, DRs would appear a well-suited application of a standardised automated detection method easily and accurately trained on a small training cohort ^19,32,33^.

The Dice similarity coefficient of 0.85 ± 0.05 achieved in our test set (ranging from 0 indicating no spatial overlap between two binary segmentation results, to 1, indicating complete overlap) would be generally categorised as ‘good to very good accuracy’, although some consideration of task complexity should be accounted for. Segmentation of a well-defined structure (such as solid organ segmentation in Computed Tomography for example in liver volumetry or radiotherapy planning) will likely achieve a higher Dice score than segmentation of complex irregular structure with poorly defined boundaries. The range of Dice scores across the test set (0.77 – 0.91) likely reflects differences in DR severity, morphological complexity or staining characteristics. Although few published studies have directly measured inter-observer variability of pathologist annotation specifically of DR regions, related liver histopathology assessments typically have showed only moderate, at best, agreement (weighted Kappa values 0.5 – 0.7), highlighting the challenge of subjective histological assessment and the potential of standardised automated detection ^34–36^. The close tracking between training and validation loss throughout training confirms the model generalises well beyond the training data and the high specificity achieved (0.98) indicates excellent discrimination against false detection — a crucial factor in histopathological analysis. The robust segmentation performance achieved from fifty-one training slides contrasts sharply to many deep learning applications which require hundreds of thousands of annotated images and offers considerable opportunities for research workflows in particular, where expert pathologist annotation time can represent a significant bottleneck. The precision (0.84) and recall (0.88) of the model were well balanced, indicating that the network is neither over-segmenting nor under-segmenting DR regions.

Regression analyses revealed KRT19 as the dominant predictor of DUCK-Net coverage (β=2.14, *p*<0.001) with SOX9 contributing additional independent variance in response to liver injury (β=0.20, *p*=0.028). The inability of α-SMA and PSR to reach significance as independent predictors despite moderate individual correlations suggests the stromal response is largely covariant with epithelial expansion, rather than contributing unique predictive information — reinforcing that DUCK-Net is primarily capturing changes driven by epithelial morphology ^37,38^. Using an IHC panel intentionally restricted to 4 core epithelial and stromal markers, the model explained 88% of DUCK-Net variance, with residual unexplained variance (12%) maximal between days 14 and 21, when DUCK-Net exceeded IHC-predicted values. This timing coincides with peak inflammatory infiltration in the DR niche, when ductular proliferation is accompanied by dense macrophage and neutrophil influx serving to promote and sustain epithelial expansion ^39,40^. This aligns with data from Wu et al. who demonstrated that Lipid-Associated Macrophage (LAM)-cholangiocyte interactions peak at maximal injury. H&E segmentation using DUCK-Net therefore offers potential to capture additional biological information not covered by this panel and unlike IHC which provides binary protein detection, offers continuous morphological information such as cellular architecture, nuclear features and tissue organisation. Such potential advantages of deep-learning based H&E analysis were further illustrated by ROC analysis which showed that DUCK-Net, unlike any single IHC marker, demonstrated excellent discrimination across multiple clinically relevant comparisons: control vs. injury (AUC 0.98, comparable to KRT19) and injury vs. recovery (AUC 0.81).

The strong alignment between DUCK-Net and the LPLC spatial transcriptomic domain described by Wu et al. despite differences in experimental design, mouse strain and quantification methods, is indicative not only of the conserved kinetics of DR biology in the DDC injury/recovery model but also the ability of H&E-based deep learning to discriminate biologically distinct subpopulations driving DR expansion and regression. Conventional Pearson correlation yielded similar coefficients for both CIRSTA domains (LPLC *r*=0.80; Chol *r* = 0.82), however DTW analysis to account for trajectory shape and magnitude revealed DUCK-Net was 3.2-fold more similar to LPLC (DTW = 0.09) than Chol (DTW = 0.30). This aligns with current understanding of DR biology informed by lineage tracing studies which have demonstrated that DRs involve activation and expansion of HPCs that subsequently develop into reactive cholangiocytes ^23,41^.

An important consideration is that in this study DUCK-Net has been tested and validated exclusively using the DDC dietary model of biliary injury. Whilst DDC produces a reliable and robust ductular reaction with characteristic periportal expansion, caution is warranted in extrapolating these findings to other contexts such as the centrilobular reactions seen in hypoxic injury or extensive HPC activation characteristic of massive hepatic necrosis. Furthermore, human liver diseases associated with DRs — PBC, PSC, metabolic dysfunction-associated liver disease (MASLD), and biliary atresia — exhibit distinct DR phenotypes defined by their unique aetiologies and future studies should evaluate performance across diverse contexts. However, the potential of DUCK-Net to offer enhanced discriminatory potential over classic IHC markers to capture the spatiotemporal dynamics of liver injury and recovery, better reflecting the molecular response of the Ductular Reaction, makes this a promising accessible research tool with few computational costs. Although our test set was purposely selected to reflect a spectrum of staining intensities, DDC response and image artefacts, and performed well during external validation, DUCK-Net can be re-trained on institution-specific data to optimise colour normalisation and generalisability. Beyond accuracy, the practical advantages for research workflows are considerable: rapid and standardised quantification of DRs otherwise requiring serial sectioning for IHC panel and/or specialist pathologist annotation. Realisation of its full potential will require systematic evaluation across the diverse liver pathologies and clinical contexts in which Ductular Reactions are observed.

## Acknowledgements

We thank Dr. Scott Waddell and Dr Elizabeth Carmichael for kindly providing H&E-stained liver sections that were ultimately not used in the final study.

## Data Availability Statement

The DUCK-Net model and analysis code are publicly available at https://github.com/WhatsMyPurpose/ducknet-dr-segmentation. Raw spatial transcriptomic data from Wu et al. were accessed from the CNGB Nucleotide Sequence Archive (accession CNP0003447) and CIRSTA database (https://db.cngb.org/stomics/cirsta/).

---

**Supplementary Figure 1.** Regression model diagnostics for multiple linear regression of DUCK-Net coverage against IHC markers.

(A) Scatterplot matrix showing pairwise relationships between DUCK-Net coverage and IHC markers (KRT19, SOX9, α-SMA, PSR). Linear relationships were observed between all variables with no evidence of non-linear patterns.

(B) Residual diagnostic plots for the multiple linear regression model. Orange lines represent LOESS smoothing curves (or theoretical normal distribution for Q-Q plot). Residuals vs Fitted (top left): random scatter indicates linearity assumption satisfied. Normal Q-Q (top right): points along the diagonal confirm approximately normally distributed residuals. Scale-Location (bottom left): relatively stable trend line indicates acceptable homogeneity of variance. Residuals vs Leverage (bottom right): no points exceed Cook’s distance thresholds, confirming absence of highly influential observations.

