## Supplementary figures and images for "DUCK-Net: Automated deep learning segmentation of Ductular Reactions in murine liver injury captures multicellular niche dynamics from H&E morphology"

### Supplemental Figure 1

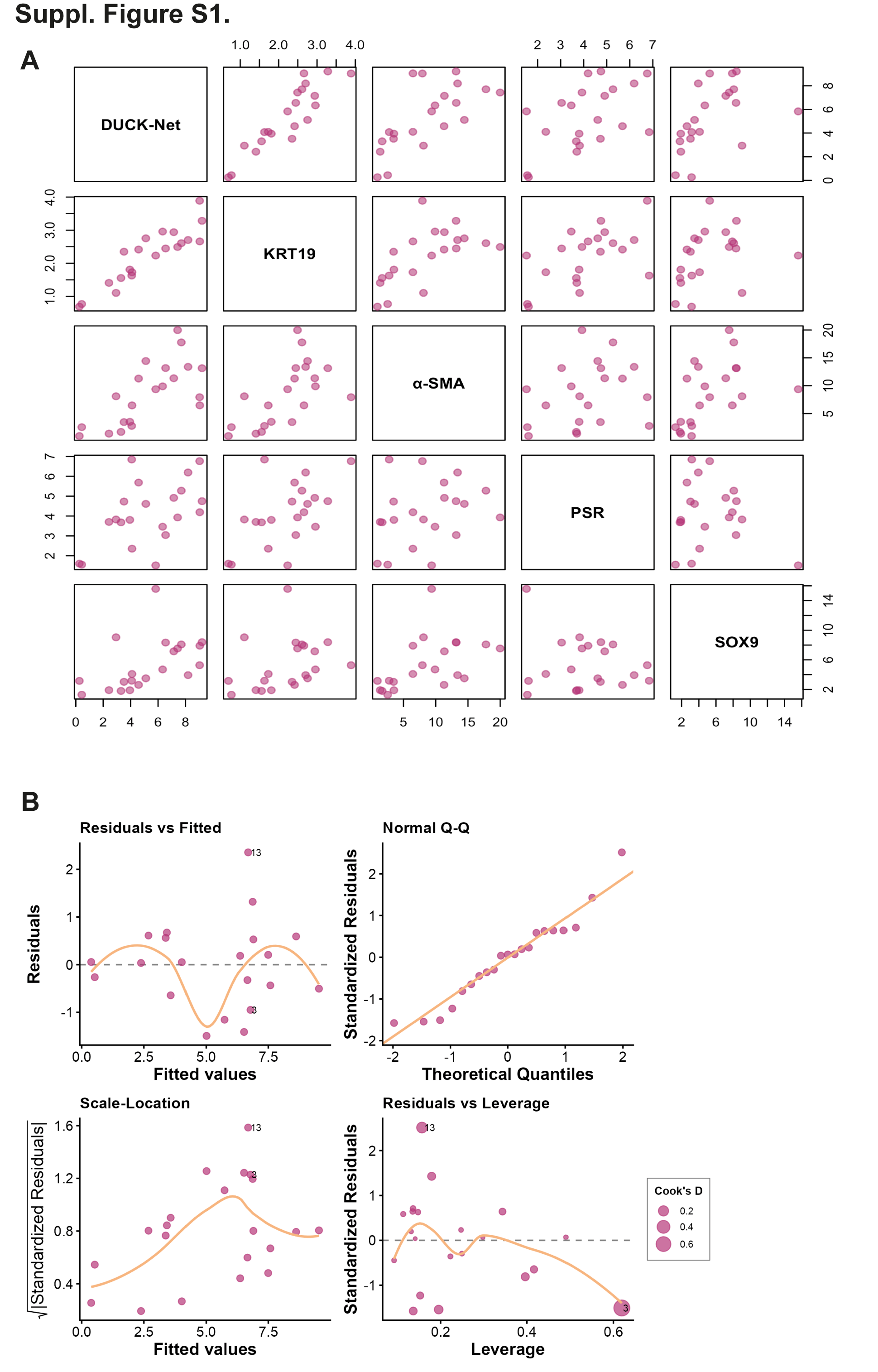
