## Supplemental Material for "DUCK-Net: Automated deep learning segmentation of Ductular Reactions in murine liver injury captures multicellular niche dynamics from H&E morphology"

**Supplementary data**

**Supplementary Methods**

**Model Architecture, Training, Validation and Testing**

Whole-slide images in NDPI format were processed using OpenSlide and down sampled by a factor of 8, providing sufficient preservation of tissue microstructure while allowing practical patch dimensions for deep learning. Pathologist annotations, exported from QuPath as JSON polygon objects, frequently encompassed vascular lumina, these were excluded through intensity thresholding followed by elliptical morphological operations via OpenCV ^20^. All image processing was implemented in Python.

The DUCK-Net model was configured with 17 initial filters. The model was trained on randomly sampled RBG patches of 512 × 512 × 3 pixels. To address the class imbalance inherent in DR segmentation, a stratified sampling strategy was implemented: sampling probability for each source image was weighted proportionally to total tissue area, and at least 50% of patches per batch were constrained to contain DR tissue. Training proceeded with a batch size of 4 and 2,000 samples per epoch. To improve generalisation and mitigate overfitting, data augmentation was implemented using the Albumentations library ^21^. Geometric augmentations including random rotations and flips were applied, along with Gaussian blur. Stain variation was addressed through a custom haematoxylin-eosin-DAB (HED) colour jitter augmentation, wherein images were transformed to HED colour space, the haematoxylin and eosin channels were independently jittered, and the image was transformed back to RGB space.

Model optimisation employed the Dice coefficient as the objective function. Learning rate adjustment followed a reduce-on-plateau strategy, monitoring the validation Dice coefficient with a reduction factor of 0.5 and patience of 10 epochs. Training was conducted on an NVIDIA RTX 6000 Ada graphics processing unit with 48 GB memory, completing in 8 hours over 88 epochs.

For validation of the model, a sliding window approach was employed with 512 x 512 pixel patches and 35% overlap between adjacent windows. Overlapping predictions were combined using a logical OR operation to produce full-resolution segmentation maps. The Dice coefficient was computed across the entire validation set, and model checkpoints were saved when validation Dice improved.

**Statistical Analyses**

Area under the ROC curves (AUC) was calculated from empirical (non-parametric) ROC curves with 95% confidence intervals estimated using the DeLong method ^22^. Optimal classification thresholds were determined by maximising Youden’s J statistic (sensitivity + specificity -1). Pairwise comparisons of AUC between DUCK-Net and each marker were performed using the DeLong test for correlated ROC curves. ROC curves were smoothed using binormal fitting for visualisation purposes only: this approach models the score distributions of each class as normal and derives a parametric ROC curve accordingly. All ROC analyses were performed in R using the *pROC* package.

**Supplementary Figure S1.**

Regression model diagnostics for multiple linear regression of DUCK-Net coverage against IHC markers.

**Supplementary Table S1.**

Annotated pixel statistics and dataset composition for model development and evaluation.

**Supplementary Table S2**.

DUCK-Net performance metrics for individual test samples.

**Supplementary Table S3.**	ROC analysis summary for DUCK-Net and IHC markers

across clinically relevant comparisons.

**Supplementary Table S4.**	Time course quantification of DUCK-Net, IHC markers, and interpolated spatial transcriptomic domains.
