## Supplemental Table 1 for "DUCK-Net: Automated deep learning segmentation of Ductular Reactions in murine liver injury captures multicellular niche dynamics from H&E morphology"

**Supplementary Table S1.** Annotated pixel statistics and dataset composition for model development and evaluation.

|  | **Annotated Pixels (Million)** | | |  |  |  |
| --- | --- | --- | --- | --- | --- | --- |
|  | Image | Tissue | DR | Image Tissue Coverage (%) | Tissue DR Coverage (%) | Samples used |
| Train | 13,535 | 8,890 | 378 | 65.7% | 4.3% | WSI  50x sample selection |
| Validation | 4,220 | 2,683 | 130 | 63.6% | 4.8% | WSI  50x sample selection |
| Test | 4,420 | 4,072 | 627 | 92.1% | 15.4% | 10x independent sample selection |
| **Total** | **22,175** | **15,645** | **1,135** | **70.5%** | **7.3%** |  |

Pixel counts (millions) are reported for total annotated image area, tissue area, and ductular reaction (DR) area, calculated from the original full-resolution whole-slide images prior to downsampling. Tissue image coverage (%) indicates the proportion of annotated image area identified as tissue, and tissue DR coverage (%) indicates the proportion of tissue area annotated as DR. Training data included one whole-slide image (WSI) split 75:25 into training and validation regions, supplemented with sampled regions from 50 additional WSIs distributed across these sets. The independent test set comprised 10 regions sampled from separate WSIs not used during training or validation.
