## Supplemental Table 2 for "DUCK-Net: Automated deep learning segmentation of Ductular Reactions in murine liver injury captures multicellular niche dynamics from H&E morphology"

**Supplementary Table S2.** DUCK-Net performance metrics for individual test samples.

| **Slide ID** | **Image** | **Dice** | **Precision** | **Recall** | **Specificity** | **Accuracy** |
| --- | --- | --- | --- | --- | --- | --- |
| 131 | 1 | 0.915 | 0.914 | 0.916 | 0.974 | 0.961 |
| 132 | 2 | 0.896 | 0.883 | 0.910 | 0.975 | 0.964 |
| 133 | 3 | 0.893 | 0.875 | 0.912 | 0.985 | 0.978 |
| 134 | 4 | 0.876 | 0.928 | 0.830 | 0.979 | 0.943 |
| 135 | 5 | 0.867 | 0.810 | 0.932 | 0.976 | 0.972 |
| 136 | 6 | 0.780 | 0.673 | 0.927 | 0.952 | 0.949 |
| 137 | 7 | 0.827 | 0.766 | 0.898 | 0.969 | 0.962 |
| 138 | 8 | 0.884 | 0.823 | 0.955 | 0.961 | 0.960 |
| 139 | 9 | 0.768 | 0.901 | 0.670 | 0.991 | 0.958 |
| 150 | 10 | 0.831 | 0.860 | 0.804 | 0.984 | 0.965 |
|  | **Mean** | **0.854** | **0.843** | **0.875** | **0.975** | **0.961** |
|  | **SD** | **±0.048** | **±0.074** | **±0.081** | **±0.011** | **±0.009** |

Performance metrics for DUCK-Net segmentation evaluated on 10 independent test samples from mouse liver sections following 21 days DDC dietary injury. Each sample was annotated by a specialist clinical pathologist. Dice coefficient (Dice), precision, recall (sensitivity), specificity, and accuracy are reported for each sample with mean ± standard deviation (SD) across the test set.
