## Supplemental Table 3 for "DUCK-Net: Automated deep learning segmentation of Ductular Reactions in murine liver injury captures multicellular niche dynamics from H&E morphology"

**Supplementary Table S3.**	ROC analysis summary for DUCK-Net and IHC markers across clinically relevant comparisons.

| **Comparison** | **Marker** | **AUC** | **95% CI** | **Optimal Threshold** | **Sensitivity** | **Specificity** |
| --- | --- | --- | --- | --- | --- | --- |
| **Control vs Any Injury (D0 vs D7-35)** | DUCK-Net | 1.00 | 1.00 – 1.00 | 1.37 | 1.00 | 1.00 |
|  | KRT19 | 1.00 | 1.00 – 1.00 | 0.94 | 1.00 | 1.00 |
|  | SOX9 | 0.95 | 0.84 – 1.00 | 1.54 | 1.00 | 0.80 |
|  | α-SMA | 0.82 | 0.65 – 1.00 | 6.23 | 0.74 | 1.00 |
|  | PSR | 0.97 | 0.90 – 1.00 | 2.26 | 0.95 | 1.00 |
| **Peak vs Early/Control (D14-21 vs D0-7)** | DUCK-Net | 0.98 | 0.94 – 1.00 | 6.08 | 1.00 | 0.89 |
|  | KRT19 | 1.00 | 1.00 – 1.00 | 2.47 | 1.00 | 1.00 |
|  | SOX9 | 0.67 | 0.36 – 0.98 | 3.55 | 1.00 | 0.56 |
|  | α-SMA | 0.90 | 0.75 – 1.00 | 9.63 | 0.86 | 0.89 |
|  | PSR | 0.98 | 0.94 – 1.00 | 3.25 | 1.00 | 0.89 |
| **Injury vs Recovery (D7-21 vs D28-35)** | DUCK-Net | 0.81 | 0.58 – 1.00 | 5.47 | 0.82 | 0.88 |
|  | KRT19 | 0.66 | 0.38 – 0.94 | 2.43 | 0.73 | 0.75 |
|  | SOX9 | 0.97 | 0.89 – 1.00 | 3.72 | 1.00 | 0.88 |
|  | α-SMA | 0.82 | 0.59 – 1.00 | 4.99 | 1.00 | 0.62 |
|  | PSR | 0.66 | 0.40 – 0.92 | 3.57 | 0.36 | 1.00 |

Receiver Operating Characteristic (ROC) analysis was performed to evaluate the discriminatory performance of DUCK-Net and conventional IHC markers (KRT19, SOX9, α-SMA, PSR) for classifying injury states in the DDC model. Area under the ROC curve (AUC) was calculated from empirical (non-parametric) ROC curves with 95% confidence intervals (CI) estimated using the DeLong method. Optimal classification thresholds were determined by maximising Youden's J statistic (sensitivity + specificity − 1). Sensitivity and specificity are reported at the optimal threshold.
