## Supplemental Table 4 for "DUCK-Net: Automated deep learning segmentation of Ductular Reactions in murine liver injury captures multicellular niche dynamics from H&E morphology"

**Supplementary Table S4.**	Time course quantification of DUCK-Net, IHC markers, and interpolated spatial transcriptomic domains.

| **Day** | **Phase** | **DUCK-Net (%)** | **KRT19 (%)** | **SOX9 (%)** | **α-SMA (%)** | **PSR (%)** | **LPLC** | **Chol** |
| --- | --- | --- | --- | --- | --- | --- | --- | --- |
| 0 | Baseline | 0.35 ± 0.09 | 0.47 ± 0.12 | 1.24 ± 0.51 | 3.82 ± 0.90 | 1.44 ± 0.28 | 0.102 | -0.026 |
| 7 | Injury | 4.86 ± 0.82 | 1.88 ± 0.30 | 9.26 ± 2.37 | 9.28 ± 1.43 | 2.68 ± 0.49 | 0.362 | -0.018 |
| 14 | Injury | 7.53 ± 0.61 | 2.92 ± 0.16 | 6.94 ± 0.79 | 13.59 ± 2.23 | 4.26 ± 0.34 | 0.514 | -0.007 |
| 21 | Injury | 8.31 ± 0.39 | 3.19 ± 0.53 | 10.90 ± 4.37 | 13.61 ± 2.56 | 5.45 ± 0.47 | 0.590 | 0.005 |
| 28 | Recovery | 5.51 ± 1.24 | 2.65 ± 0.48 | 3.30 ± 0.74 | 8.84 ± 2.73 | 5.18 ± 0.67 | 0.626 | 0.000 |
| 35 | Recovery | 3.49 ± 0.38 | 1.80 ± 0.20 | 2.50 ± 0.35 | 2.79 ± 0.49 | 4.77 ± 0.73 | 0.518 | -0.008 |

Quantification of DUCK-Net coverage and IHC markers across the DDC injury and recovery time course. DUCK-Net coverage and IHC markers (KRT19, SOX9, α-SMA) are expressed as % positive area or cells; PSR (picrosirius red) as % collagen area. Values shown as mean ± SE (n = 4–5 per timepoint). LPLC (Liver Progenitor Like Cell) and Chol (Cholangiocyte) domain scores were derived from spatial transcriptomic (Stereo-seq) data from Wu et al. (2024) and interpolated to match experimental timepoints using linear interpolation.
